# Bayesian linear models with unknown design over finite alphabets

**DOI:** 10.1101/2022.10.20.513021

**Authors:** Yuexuan Wang, Andreas Futschik, Ritabrata Dutta

## Abstract

Our topic is the reconstruction of the unknown matrices *S* and *ω* for the multivariate linear model *Y* = *Sω* + *ε* under the assumption that the entries of *S* are drawn from the finite alphabet 𝔄 = 0, 1 and *ω* is a weight matrix. While a frequentist method has recently been proposed for this purpose, a Bayesian approach seems also desirable. We therefore provide a new hierarchical Bayesian method for this inferential task. Our approach provides estimates of the posterior that may be used to quantify uncertainty. Since matching permutations in both *S* and *ω* lead to the same reconstruction *Sω*, we introduce an order-preserving shrinkage prior to establish identifiability with respect to permutations.

## 1. Introduction

### Linear models with unknown design over finite alphabets

Linear models are among the most widely used statistical models, where we consider a matrix *Y* of observations, and its factorization into the product of a design matrix *S* and a matrix *ω* of parameters of interest. More specifically, under some assumptions on the noise (e.g. Gaussian noise) we infer *ω* given a known design matrix *S*.

Here we address the reconstruction of both *S* and *ω* under the assumption *Y* ∈ [0, 1]^*N*×*T*^ and the design matrix having entries in {0, 1}. Since we want to reconstruct two unknown matrices from their product, they are not identifiable and some conditions on the parameter matrix will be needed. In this work, we aim to provide a Bayesian method to accurately reconstruct *S* and *ω* jointly. We will provide a proper way to model the noise as well as quantify the required constraints in the sense of Bayesian hierarchical modeling, and introduce an efficient sampling scheme to obtain estimates for both matrices. In Section 2.1 we explain our model and provide details about the identifiability conditions.

Different approaches exist for reconstructing the design and parameter matrices from the observed data. We will give a review below regarding to the existing methodologies in different fields of study.

Linear models with unknown finite alphabet designs has been used successfully in genetics. More specifically, it has been used for the purpose of haplotype reconstruction from population genetic data. A haplotype is a segment of DNA that is inherited from a single parent. Haplotypes carry crucial genetic information, and their reconstruction is therefore desirable in many situations. Inferring haplotypes from diploid genotype data is called phasing, and considerable efforts have been taken to develop methodology for this purpose [1, 2, 3]. In population genetic applications, however, individuals are often sequenced in pools so that individual genotype information is unavailable. This is to save cost in terms of time and money [4, 5, 6]. Such pool sequencing data provide allele frequencies at a single-nucleotide polymorphism (SNP) level (*Y* ∈ [0, 1]^*N*×*T*^) [7, 8] rather than haplotype information. In such an application, *N* represents the number of considered SNPs, and *T* will be the number of time points at which sequencing information is available. The goal is then to infer the haplotype structure *S* ∈ {0, 1}^*N*×*m*^ and the haplotype frequencies at different time points *ω* ∈ [0, 1]^*m*×*T*^ from the allele frequency matrix *Y*. In the context of Evolve and Resequencing (E&R) experiments, [9] introduced a method that reconstructs only one selected haplotype from replicated Pool sequencing temporal data. There are also some approaches aimed to reconstruct haplotype frequencies with the corresponding founder haplotypes known [10, 11, 12, 13, 14]. [15] proposed an adaptation of the work by [16] to reconstruct both haplotype structure and haplotype frequency without prior information. Here, we will provide a Bayesian method that can be applied to such problems.

Another relevant application of this model can be found in the field of digital communication, where the elements of the signal waveform matrix *S* are assumed to belong to a finite alphabet, where *N* represents the number of symbols per data burst and *m* is the number of signals. *ω* is the array response matrix with *T* representing the number of sensors. [17, 18, 19] prove that bilinear decomposition is unique under constant modulus (CM) and finite alphabet (FA) constraint. To separate the spatial response and signal waveform from synchronous co-channel digital signals, a maximum-likelihood approach is proposed by [17]. [20] incorporate fractional sampling and blind source separation (BSS) to recover transmitted symbols with inter-symbol interference.

Bayesian penalization has been well-studied for years [21, 22, 23, 24, 25, 26]. Instead of putting a penalty term under the Frequentist paradigm, we introduce a prior distribution to penalize the effects in a Bayesian way. It helps us to shrink insignificant effects to zero and perform model selection as well as provide more flexibility to various penalties that serve our needs. The penalization in MCMC sampling is addressed by a specific prior on the corresponding parameter, which is called shrinkage-prior. For non-convex penalties, MCMC has the advantage of easy implementation over optimizing objective functions. There are two groups of shrinkage prior: spike-and-slab priors [27, 28, 29, 30] referring to a mutually independent relationship among effects with a mixture distribution consisting of spike (degenerate distribution at zero) and slab (diffused distribution) and the global-local shrinkage prior [31, 32, 33]. The second group of priors is more computationally efficient than the spike-and-slab prior and has better mixing properties. It has two parts of shrinkage; a global parameter controls overall shrinkage while the local parameters control effect-specific shrinkage.

### Our contribution

In this paper, we propose a Bayesian approach to solve the unknown design over finite alphabets problem, which has not yet been solved in a Bayesian fashion. Inspired by [34], we construct a novel order-preserving shrinkage prior to impose a decreasing (in norm) order of the rows of the parameter matrix *ω*. We addressee this reconstruction problem under a Bayesian framework, which allows us to quantify estimation uncertainty. The details of the algorithm will be fully discussed and its performance will be investigated. A comparison with existing methods will also be provided. The paper is organized as follows: in Section 2, we propose a hierarchical Bayesian model to reconstruct the target matrix under the constraints we specified. In Section 3.1, experiments with a set of simulated data are examined and compared with an existing frequentist method. Section 4 investigates the performance of our method on an HIV data set.

## 2. Bayesian linear model for finite alphabet design

### 2.1. Model specification

We consider the following linear model with Gaussian error:

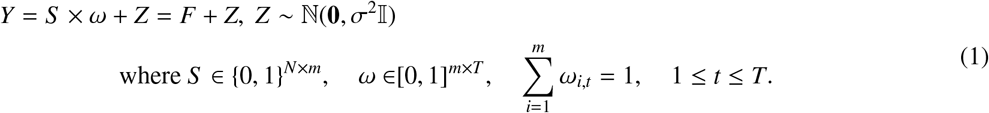

We denote Θ_*S*_, Θ_*ω*_ and Θ_*F*_ to be the parameter spaces of *S, ω* and *F* respectively satisfying the constraints provided in Equation 1. In particular, the entries of *S* come from a finite alphabet 𝔄 = {0, 1} and entries for *ω* and *F* should be between 0 and 1.

#### Identifiability of the solution

The recovery of *S* and *ω* matrix is non-identifiable without proper constraints. Hence, in [35], the authors stated some unidentifiable situations: (i) If we do not fix the ordering of parameter matrices, for any permutation matrix *A*, we will have *Sω* = *S AA*^−1^*ω*, hence we get the same reconstruction when both *S* and *ω* are permuted analogously. (ii) For the alphabet {0, 1}, when we have two or more identical rows in *ω*, the corresponding entries in S are not unique. (iii) If there exist two or more identical columns in *S*, suitable changes of the corresponding weights lead to the same reconstruction. For more details on identifiability issues, see [35].

We deal only with (i) in our modelling, since situations (ii) and (iii) occur on subsets of the parameter space that have either probability zero, or a very low prior probability. To resolve situation (i), we propose an order preserving prior to favour matrices *ω* where the rows are ordered by their norms.

### 2.2. Bayesian Hierarchical Model and Gibbs Sampler

There have been many Bayesian approaches developed over the last few decades to serve the purpose of drawing samples from the posterior distribution. The simplest situation is that the integral required for a posterior can be directly evaluated by numerical integration. Unfortunately, this approach is limited to low-dimensional problems. For situations where target distribution is difficult to sample from or simply intractable, advanced Bayesian approaches are needed. One classical approach of the Monte Carlo technique is the rejection sampling. It draws samples from a target probability density function by accepting or rejecting samples from a known proposal distribution with an acceptance probability, which is calculated by the ratio of the densities. The proposal distribution here is a known density that can cover the target distribution by multiplying a scalar. This bound for the acceptance ratio is unable to be established for complicated target. To generate samples from more complicated target distribution, Monte Carlo Markov chain (MCMC) technique is preferred. But currently, the popular gradient based MCMC schemes (e.g., Hamiltonian Monte Carlo) can not be used for discrete parameter space as gradients can not be computed in this case [36, 37]. Hence, they can not be used in our situation as our sample space consists of the design matrix *S* which takes values from the finite alphabet {0, 1}, making many of the popular MCMC packages unusable for our problem. Hence, we developed a blocked Gibbs sampling scheme, which is described in the rest of this section.

#### 2.2.1. Prior Specification

We adopt a Bayesian approach and assume independent hierarchical priors on the unknown parameter matrices *S, ω* and the noise *σ*^2^. Here we introduce details of all prior choices.

##### Prior on binary matrix S

For the binary matrix *S*, we assume a Bernoulli prior. More specifically, we introduce Bernoulli priors with row specific hyperparameters, i.e. *S* _*i,k*_ ∼ Bernoulli(*p*_*i*_), *i* ∈ [*N*]. And we assume *p*_*i*_ to be independent with beta conjugate prior with hyperparameters *a*_*β*_ and *b*_*β*_.

##### Prior on ω

In order to satisfy the identifiability condition we described in Section 2.1, it is important that the prior on *ω* guarantees a unique ordering of the rows of *ω*. We therefore adapt a shrinkage prior proposed by [34] to introduce decreasing row sums for the *ω* matrix. The prior specification below, introduces a degree of shrinkage that increases with the row index.

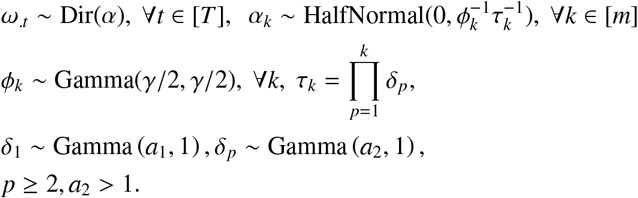

In this shrinkage prior, *τ*_*k*_ is a global shrinkage parameter for the *k*^*th*^ row, and *ϕ*_*k*_ is a local shrinkage parameter. *γ* is a self-defined hyperparameter for *ϕ*, we choose 3 in our experiments. *a*_1_, *a*_2_ are self-defined hyperparameters with the lower bound 0 and 1 respectively since they are hyperparameters of the Gamma distribution and *a*_2_ should greater than 1 to guarantee *δ*_*p*_ has expectation larger than 1. Then 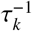 are stochastically decreasing in *k*. Hence, we impose the decreasing order of rows in *ω* by introducing decreasing variance to *α*_*k*_. However, if we only include *τ*_*k*_ in the variance of *α*, it will have a tendency to over-shrink the nonzero rows of *ω*. Hence the local shrinkage parameter *ϕ*_*k*_ is introduced.

##### Prior on σ ∈ ℝ^+^

We consider an Inverse Gamma distribution as prior on *σ*, which is the noise term of our model. Hence *σ* ∼ InverseGamma(*a*_*ξ*_, *b*_*ξ*_), where *a*_*ξ*_ and *b*_*ξ*_ are hyperparameters (we both choose 3 in our experiments).

#### 2.2.2. Posterior computation for finite alphabet design Initialization of parameters

- *ω*^*init*^: The initialization of *ω* is obtained using the method provided by [15], the details of this algorithm can be found in the supplementary information (algorithm 2).
- *S* ^*init*^: for each row of *S* ^*init*^, 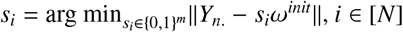
- (*σ*^2^)^*init*^: var(*Y*^*obs*^ − *S* ^*init*^ *ω*^*init*^)
- 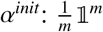.
- For all other hyperparameters, we draw them from the prior.

##### Gibbs Sampling

To sample *S, ω* and all hyper-parameters with the self-defined number of *m* from the joint posterior, the Gibbs sampler[38] is introduced. It provides a computationally efficient way to update multiple parameters of interest by blocking, and accelerates mixing with the shrinkage prior on *ω*. The hierarchical structure of our model is displayed in Figure 1):

**Figure 1:**
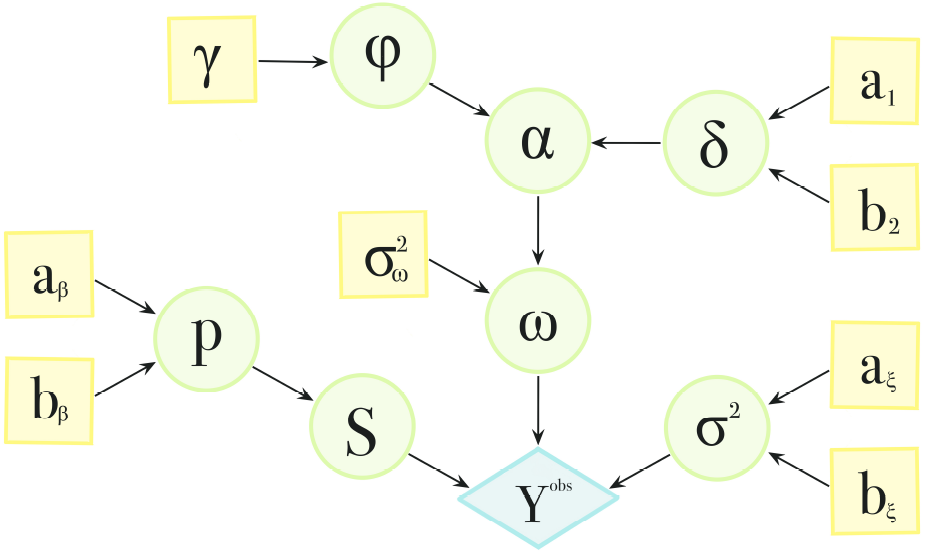
Illustration of the hierarchical structure of parameters and hyper-parameters for the blocked Gibbs sampler. The blue circle represents observed data. Green circles represent parameters to be estimated, contain our target matrices *S* and *ω*, and yellow squares indicate hyperparameters.

Our sampler proceeds as explained below.

*Step* 1 Sample *S* ∈ {0, 1}^*N*×*m*^ from the distribution:

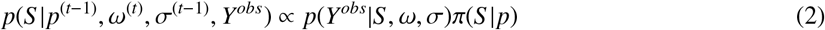

The exact sampling scheme for *S* is shown in Equation (10) below.

*Step* 2 Sample *p*_*i*_, *i* ∈ [*N*] from conjugate posteriors:

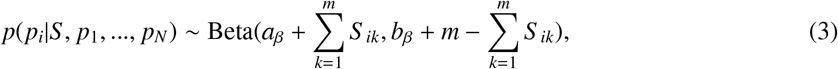

where *a*_*β*_ and *b*_*β*_ are hyperparameters (we both choose 2 in our experiments) for the prior of *p*_*i*_, i.e. *p*_*i*_ ∼ Beta(*a*_*β*_, *b*_*β*_).

*Step* 3 Sample the posterior *ω* by Metropolis-Hastings from the following joint posterior distribution with the reparametrisation illustrated below.

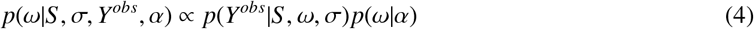

*Step* 4 Sample *α* from its posterior:

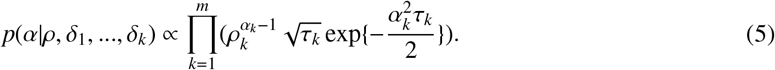

where *ρ* is the row sum of *ω* matrix divided by the sum of *ω* matrix.

*Step* 5 Sample *δ*_*k*_, *k* ∈ [*m*] from conjugate posterior:

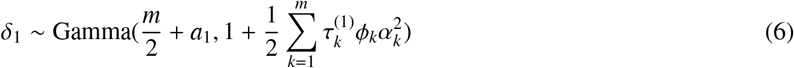

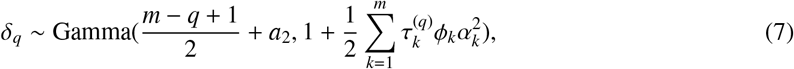

where 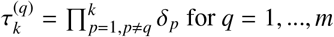.

*Step* 6 Sample *ϕ* from it’s conjugate posterior:

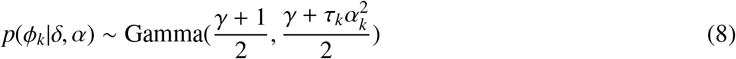

where 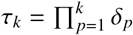 and *γ* is a hyperparameter.

*Step* 7 Sample *σ* from conjugate posterior:

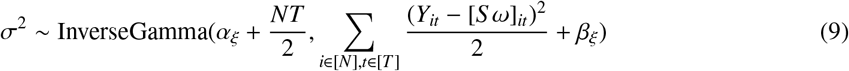

##### Exact sampling of the rows of S

We are going to update the S matrix row by row. More specifically, for the *i*^*th*^ row of the S matrix, with all other entries of S fixed, the probability mass function will be proportional to

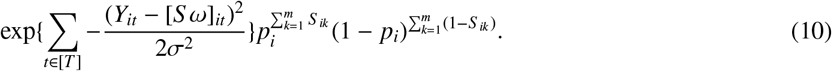

Suppose the S matrix has m columns. Then for each row, there are 2^*m*^ possible combinations of entries. One of these combinations is chosen according a the probability mass function proportional to equation(10). We are able to draw the rows of the S matrix with the corresponding probability.

##### Reparametrisation to sample ω

Since *ω* is constrained, sampling such parameters causes the problem that samples might fall outside of the domain. This will lead to inefficient sampling with a low acceptance rate. Hence, to encourage better mixing of the sampling of *ω*, we propose the following reparametrization during the sampling, which is then transformed back to obtain the actual draws. Notice that in our case, we assume that the columns of *ω* matrix are independent. To start with, we transform each column of the *ω* matrix from [0, 1] to (−∞ +∞) by the stick-breaking transformation described in the equation 11. After transforming all columns, we denote the matrix as *W*. Then we add a Gaussian error with variance 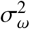 (a self-defined hyper-parameter, and we chose 0.01 in our experiments) to *W*, and this perturbed matrix is denoted as *W*^*e*^. Then the candidate *ω*^*cand*^ can be obtained by applying inverse stick-breaking transformation on *W*^*e*^, details are provided in the equation 12. Further information of the stick breaking can be found in [39].

**Forward** Suppose *ω*_.*t*_ = (*ω*_1*t*_, *ω*_2*t*_, …, *ω*_*mt*_), where *ω*_*kt*_ ∈ (0, 1), ∀*k* ∈ [*m*], ∀*t* ∈ [*T*] and 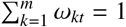.

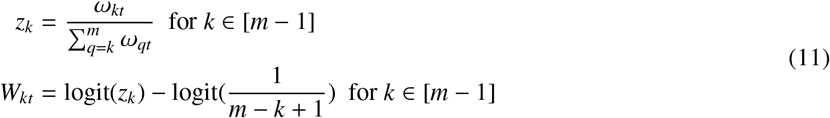

**Backward** Suppose 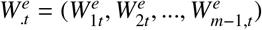, where 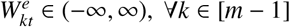

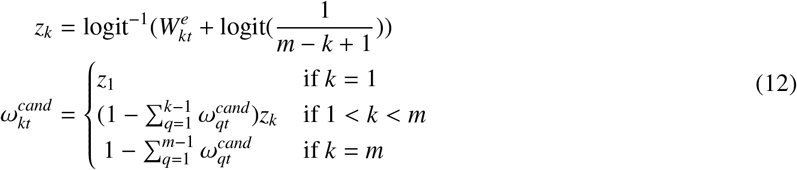

We explain the backward step first, which transforms a vector 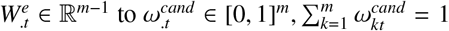. It starts by taking a stick with unit length, and for each step in the stick-breaking process, a piece of the stick is broken off and labeled with the length of the break ratio (*z*_*i*_) multiplied by remaining length of the stick 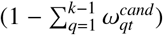 iteratively. The stick ratio is taken by the inverse logit transformation with an adjustment term logit(1/(*m − k* + 1)) to avoid transforming a zero vector to a non-uniform vector. Then the forward step can be easily attained by taking the inverse of the backward transformation.

**Absolute Jacobian determinant** To obtain the distribution of *ω* after change of variable, we need the Jacobian determinant. The Jacobian *J* is lower-triangular matrix, and upper entries are:

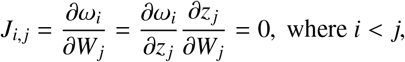

because in the stick-breaking process, the length we break off at *i*^*th*^ step (*ω*_*i*_) is independent of the proportion that will be broken off in the future step (*z* _*j*_, *i* < *j*). And the diagonal entries of the Jacobian of the inverse transformation are:

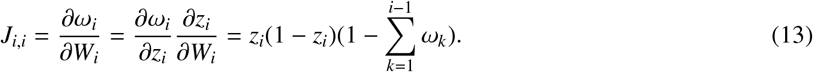

Since the Jacobian is a diagonal matrix, only the product of the diagonal entries remains in the calculation of determinant. The absolute Jacobian determinant is computed as follows:

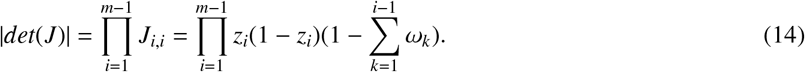

## 3. Simulation Studies

### 3.1. Randomly generated S and ω

To illustrate our method, we run simulation experiments over a range of model parameters. In particular, we chose our model dimensions within *N* = {100, 200, 500}, *m* = {3, 4, 5} and *T* = {10, 15, 20}. We simulated each entry of the S matrix independently from a Bernoulli distribution with parameter *p* = 0.5 and each column of the *ω* matrix independently from a Dirichlet distribution with parameter 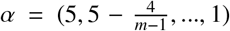. In order to obtain *Y*, we add Gaussian noise with standard deviation 0.02 to *F* = *Sω*. For each combination of these parameters, we simulate 20 data sets to test our algorithm. We report the accuracy rate and the MSE of the posterior mean estimate of S and W in comparison to the true value in Figure 2. Looking at each sub-figure, we can tell that the estimates of *S* and *ω* are improved when *T* increases. Furthermore, if we compare boxplots for the same *m* and *T* bit different *N*, it can be seen that increasing the number of rows in *Y* makes the performance better. An increase of the inner dimension parameter *m* negatively influences the estimation performance. Additional results for Gaussian noise with standard deviation 0.05 are reported in Figure A.8 in the Appendix, which confirm the findings from Figure 2. Further, we compared our posterior mean estimates with those obtained by the frequentist method from [15]. We found the accuracy of both approaches to be very close. Indeed, the mean (and variance) difference in the accuracies for S were 2e-02 (2e-03). The differences between the obtained MSE values for W are 5e-04 on average (variance 4.81e-06).

**Figure 2:**
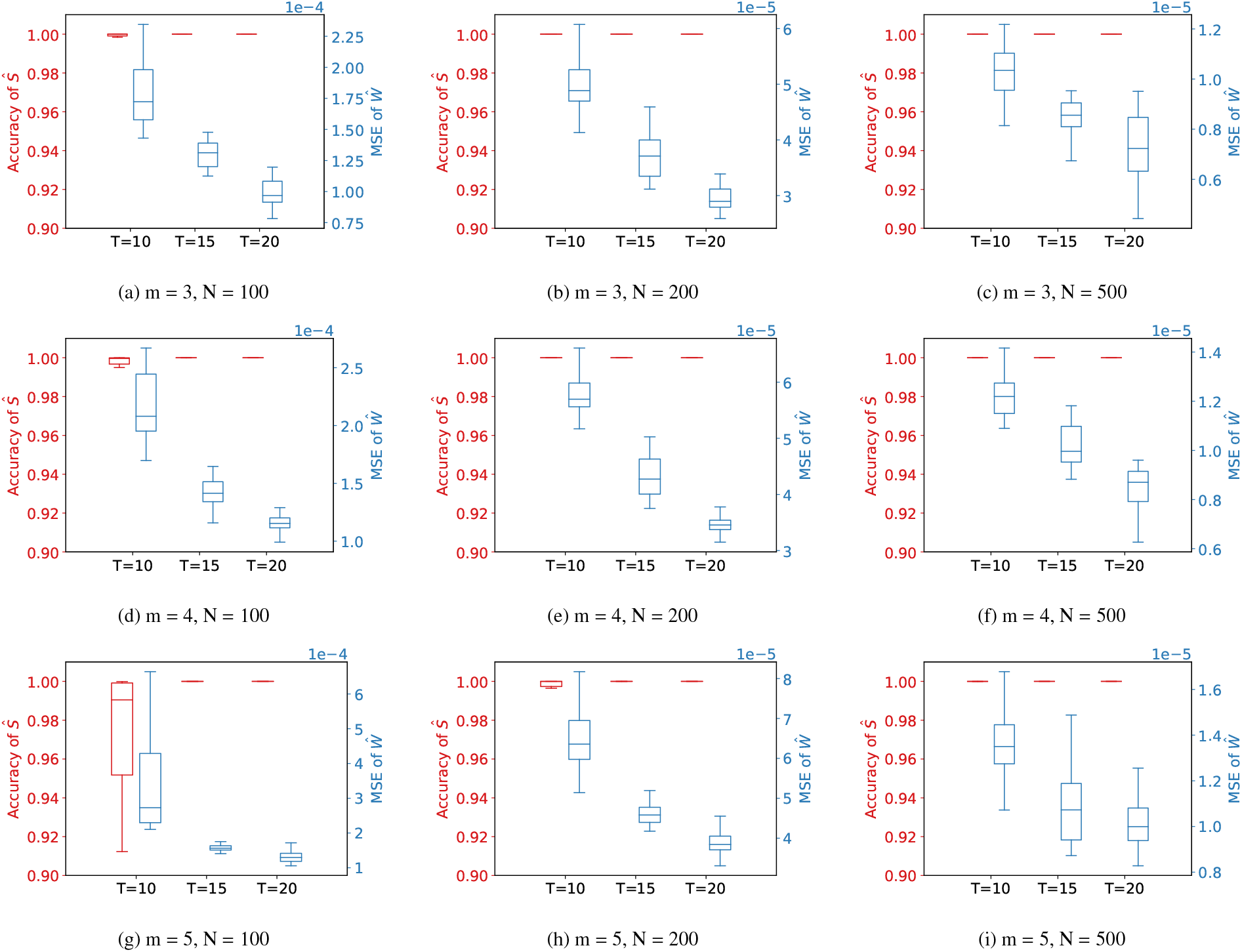
Performance of our estimates for different numbers m of haplotypes, numbers N of SNPs and numbers T of time points. Our noise level is *σ* = 0.02. The red boxplots present the accuracy rates of the estimated *S* matrix. These rates are calculated by the percentage of entries correctly estimated with the estimate being either 0 or 1 depending on which one has the higher posterior probability. The blue boxplots summarize the MSE between the true matrix *ω* and its posterior mean estimate. Y-axes are in the corresponding color. Each boxplot summarizes the results from 20 simulation runs. The estimates are obtained using the posterior based on 3000 steps of MCMC sampling excluding the burn-in steps.

To illustrate the uncertainty quantification provided by our posterior, we provide the difference of the posterior mean from the true *S* in Figure 3 for a typical simulation run. Furthermore, credible intervals for *ω* are also provided. This illustrates the uncertainty information that can be obtained in our context.

**Figure 3:**
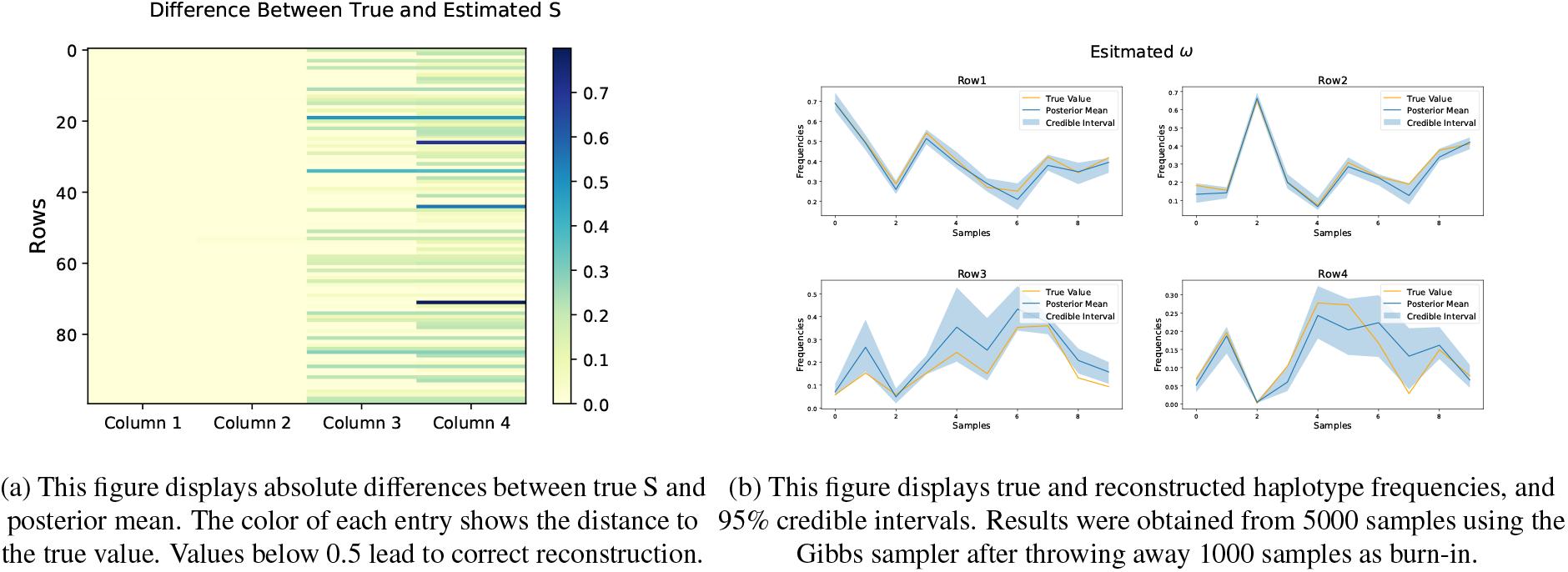
These plots are the result of simulation involving 4 haplotypes, *Y* observed under *σ* = 0.1 level of noise to investigate the performance of the credible interval.

### 3.2. Multicollinearity in S

When inferring haplotypes in a population genetic scenario, we usually face the problem that the underlying haplotypes are correlated with each other. It is well known that the variance of estimates in linear models becomes larger with increasing multicollinearity of the predictors. The same phenomenon applies to our setup. We therefore investigate, if our method is accurate enough for typical haplolotype structures encountered in population genetic data. For this purpose, we consider genome sequences that represent 202 isofemale lines of *Drosophila simulans* from an experiment described in [4]. From this data set, we randomly draw 4 founder haplotypes and focus on a 100 SNP region from position 363 to 49337 on chromosome 2L to construct our *S* matrix. And for *ω*, simulate an evolve and resequence experiment over 150 generations assuming one beneficial SNP with a selection coefficient *s* = 0.05. Multinomial sampling is used to mimic genetic drift. The population size is 500 and assumed to be constant over 150 generations, and we observed the allele frequencies every 10 generations.

In Figure 4, we provide the results of a simulation run under the described scenario. It turns out that haplotype 4 is not reconstructed well, and at some time-points the frequencies got mixed up. We attribute this result to the multicollinearity in the matrix *S* which makes estimation more difficult. To further analyze this situation, we considered different levels of correlation between the columns of *S* by replacing varying proportions of SNPs in the haplotypes by i.i.d. Bernoulli entries. This leads to scenarios with different amounts of correlation. Simulation results under these scenarios can be found in Figure 5. As can be seen, the performance of the reconstruction decreases with increasing correlation level.

**Figure 4:**
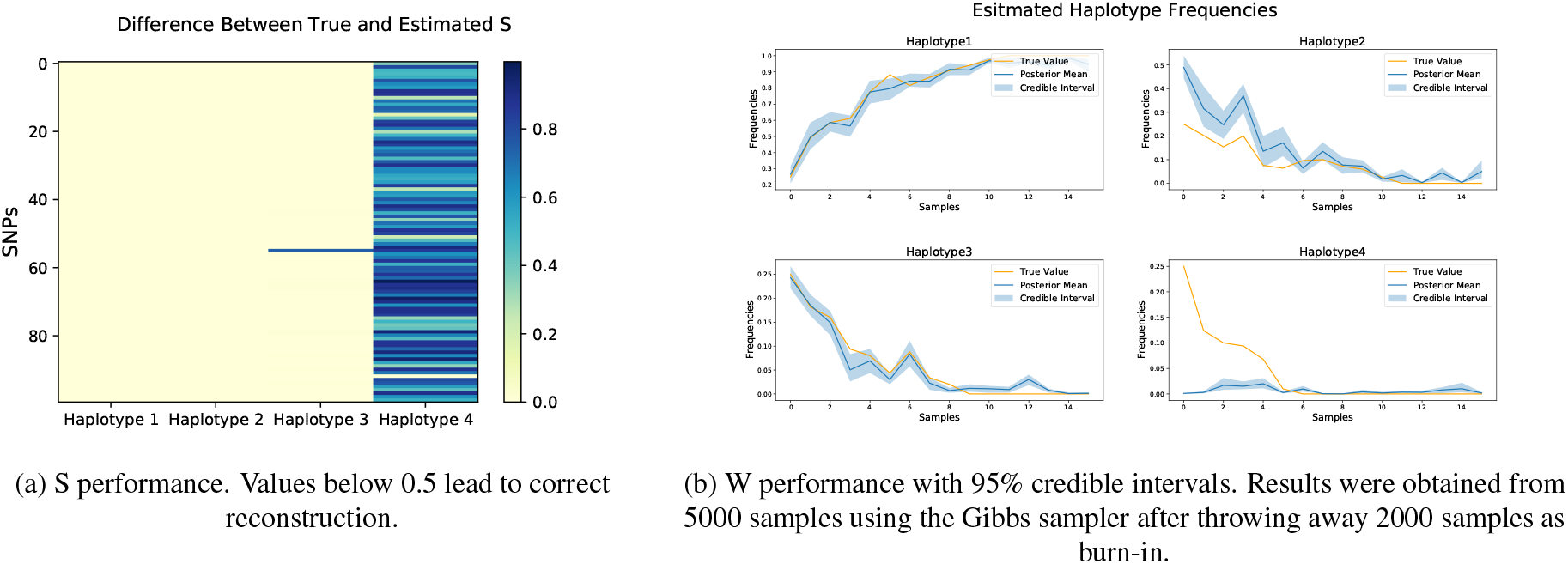
These plots are the result of simulation involving 4 haplotypes randomly drawn from Drosophila simulans data, *Y* observed under *σ* = 0.05 level of noise.

**Figure 5:**
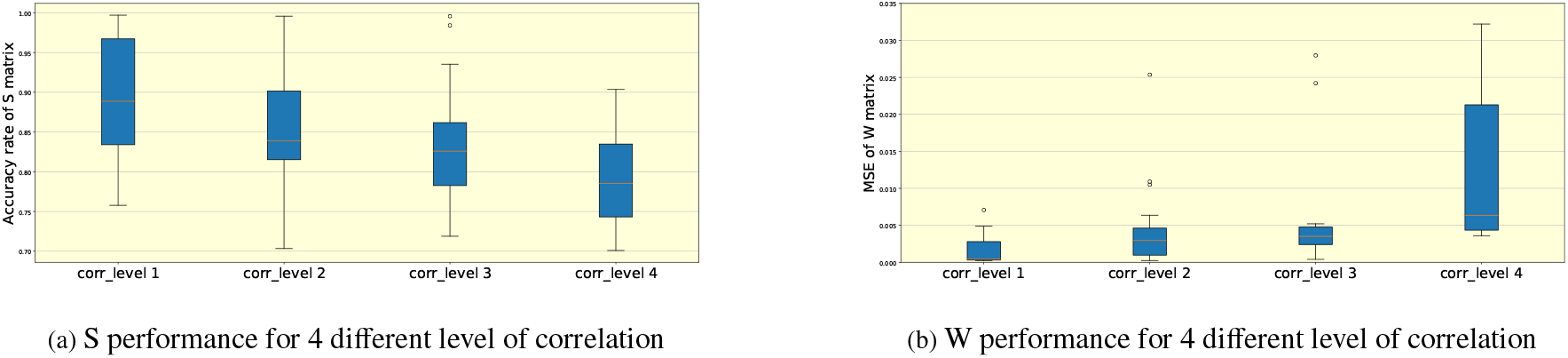
In this figure, we show the reconstruction result under 4 different levels of correlation (0.11, 0.24, 0.32, 0.43) in haplotype structure. For level 4, we use the *m* = 4, *N* = 100 case from the above simulation. At level 1-3, we randomly replace 75%, 50%, and 25% percentage of SNPs in S by Bernoulli sampling. Plot(a) is the accuracy rate of the estimated S matrix under different levels of correlation. It has a strong tendency that the accuracy rate decreases with the increase of correlation in the data. Plot(b) shows the MSE of the estimated *ω* matrix, where we can conclude the same result as figure(a).

## 4. Real data application

We applied our method on two real data sets, one being the longitudinal HIV data set provided by [40] and the other being from *Caenorhabditis elegans* (*C. elegans*) provided by [41].

### 4.1. HIV data

From the data set provided by [40], we took the data of patient 3, who was infected with HIV-1 type B. For this patient, data from 10 time points between 146 days after infection and 3079 days are available. After excluding the time points 1126, 1934, and 2927 days, which contain lots of missing values, 7 time points remain in the data. The high mutation rate of HIV leads to the constant appearance of new SNPs and haplotypes over time. Our method will therefore not be able to reconstruct all haplotypes, but rather the dominant ones. The original data contains 9095 base pairs (bp). Here we reconstruct haplotypes for a 1000 bp window that also contains the ‘vpu’ region (246 bp). For this region true genomes for 144 haplotypes are available from [40] that help us to evaluate the accuracy of our estimation results.

The results shown in Figure 6(a) indicate that the dominating haplotypes can be reconstructed very accurately. Indeed, the estimated haplotype matrix has only 0.8% mismatch. As shown in Figure 6(b), *ω* is also estimated quite accurately. The systematic deviations in the estimated frequencies are caused by the model misspecification. As we reconstruct only four out of 144 haplotypes, our estimates are bound to overestimate the true frequencies as our estimates require that the frequencies of the reconstructed haplotypes add up to one. This is in particular true, if the remaining haplotypes contribute a non-negligible proportion of the overall frequency. Unfortunately the number ov available samples does not permit to estimate the full set of haplotypes. This applies in particular to low frequency haplotypes that can usually be considered to be less important for the infection dynamics.

**Figure 6:**
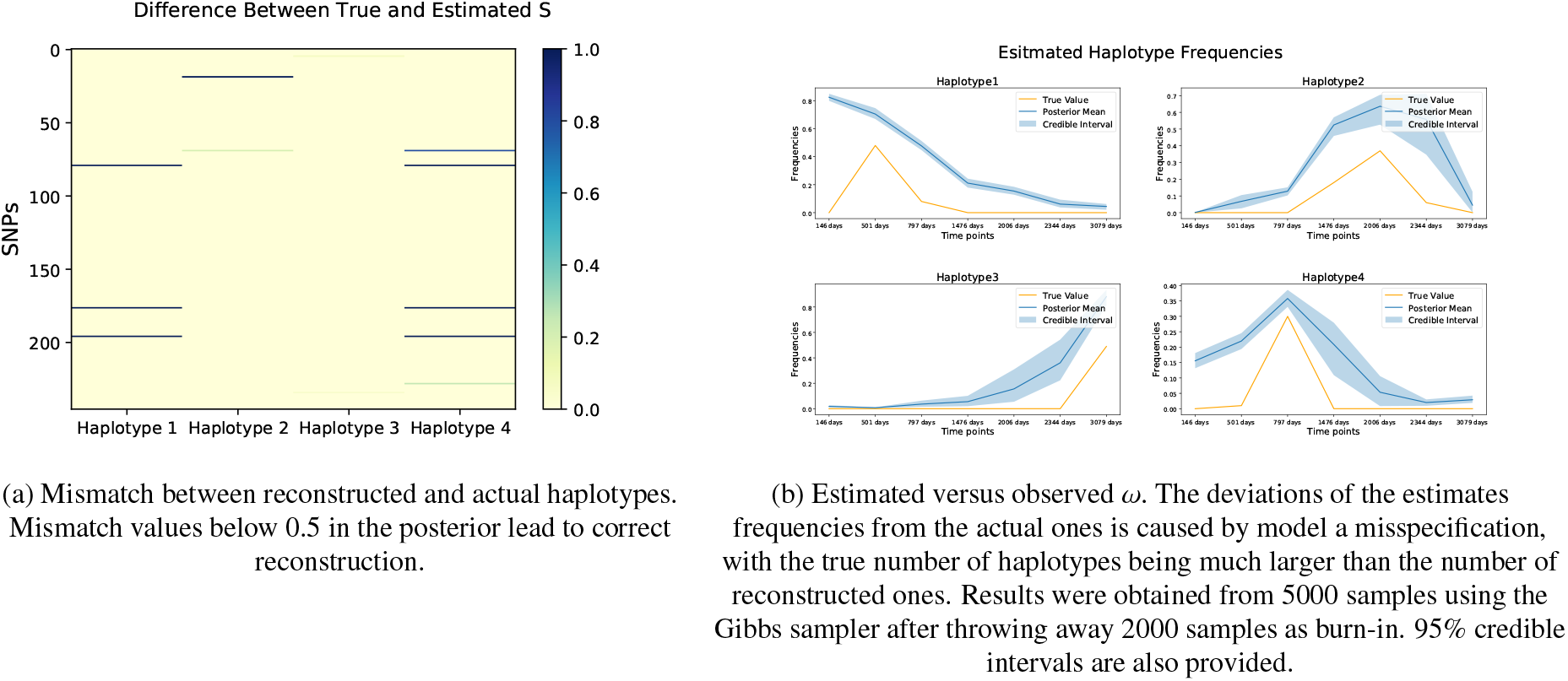
This figure provides results on the reconstruction accuracy for region ‘vpu’ of the HIV virus material sampled from patient 3 in [40].

### 4.2. Caenorhabditis elegans (C.elegans) data

In [41], an experiment concerning the evolution of C. elegans in an environment with increasing quantities of NaCl has been conducted. Data for three replicate populations have been obtained at times between generation numbers 50 and 100 under variable sex ratios and breeding modes. The census population size is 10^4^, and the effective population size is around 10^3^.

By reconstructing the haplotype structure from this data, we benefit from the availability of replicate populations that share haplotypes. We applied our reconstruction method in the genomic region 14924777-15216613 bp on chromosome 5, which contains 666 SNPs. The reconstructed haplotype structure for this segment is provided in Figure 7, where we take the most similar sequenced founder haplotype as reference. Unfortunately, no true haplotype frequency data are available for this experiment.

**Figure 7:**
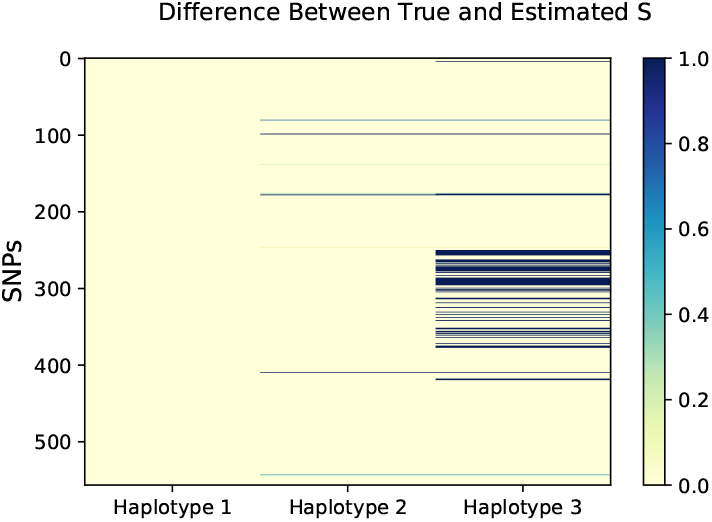
Difference between the estimated haplotype matrix *S* and the corresponding most similar sequenced founder haplotypes for our considered

Figure 7 shows the result of the C. elegans population with replicates is consistent with the data set we examined before, where haplotypes with higher frequencies perform better in the reconstruction. As a consequence, our method works well for these two real datasets, which contain both single and multiple replications.

## 5. Discussion and Conclusion

The joint recovery of both an unknown design and parameter matrix from noisy observations in a linear measurement system without prior knowledge is a challenging problem. As in the linear inverse problem, we usually deal with only one unknown matrix rather than two. Under the condition that the design matrix can only attain values from the finite alphabet 𝔄 = {0, 1}, this problem can be solved, and applied to many fields like population genetics and digital communication. In a frequentist setup, this problem has been considered in [15] and [16]. In this work, we discuss how we can recover both matrices by using Bayesian hierarchical modelling and MCMC sampling. When both of the matrices are unknown, an identifiability problem arises. To prevent an infinite number of equally good reconstructions, we imposed weak identifiability on the parameter matrices. In the Bayesian paradigm, these constraints can be modelled by carefully designed priors. Since the parameters of interest are both matrices, common MCMC methods are not applicable, especially since *S* is a binary matrix, and the entries of *ω* are bounded. For *S*, we sample the matrix row by row. And for *ω*, a stick-breaking reparametrization was applied to extend the domain of entries to (−∞, ∞). The design of the sampler encourages better mixing and efficient sampling.

Our simulation studies suggest that our algorithm provides accurate reconstructions, comparable in quality to the minimax method provided by [15]. Furthermore, our Bayesian method can quantify the uncertainty surrounding the estimation, going beyond the point estimates given by the frequentist method. Throughout the experiments, it was discovered that the shape parameters had a significant impact on the reconstruction performance. Furthermore more rows or columns in the observed matrix provide more information to improve reconstruction. However, a larger inner dimension parameter might lead to suboptimal performance. Additionally, a highly correlated true *S* matrix would lead to over-shrinkage of the estimate of *ω*.

Further, we illustrate how our method can be applied in the context of population genetics to infer haplotypes that can provide important insights into genomic phenomena. Often only allele frequency data are available and therefore haplotypes must be inferred. We looked at two real data sets, one with single (HIV) and one with multiple replications (C. elegans). Due to the restriction that the columns of *ω* sum up to one, the haplotype frequencies get overestimated, if the number of haplotypes is too large for all of them to be estimated, and non-reconstructed haplotypes are sufficiently abundant. Nevertheless our Bayesian approach can reconstruct the dominant haplotypes also in a high-dimensional setting.

To our knowledge, our work provides the first Bayesian approach to jointly reconstruct both design and parameter matrices in a multivariate regression setup with binary design. Compared to previously proposed frequentist methods, we also provide uncertainty quantification based on the posterior. We illustrated that our estimates are very accurate under a range of scenarios and also explored limitations in high dimensional settings. Besides the practical applications provided, we believe our work could be useful in a variety of practical fields. Challenges to be addressed in future work include ways to better cope with correlated design matrices. Another future direction could be to develop an algorithm to automatically select the inner dimension parameter *m* during the MCMC sampling. One might also want to specify the response in terms of a generalized linear model in order to enrich the set of available models.

## Data and Code Availability

The data mentioned above and the corresponding code are available at: https://github.com/Yuexuanwang/Bayesian-reconstruction

## Acknowledgement

We are grateful to Marta Pelizzola for her advice and help with the real data applications. RD is funded by EPSRC (grant nos. EP/V025899/1, EP/T017112/1) and NERC (grant no. NE/T00973X/1).

## Appendix A. Appendix

**Figure A.8:**
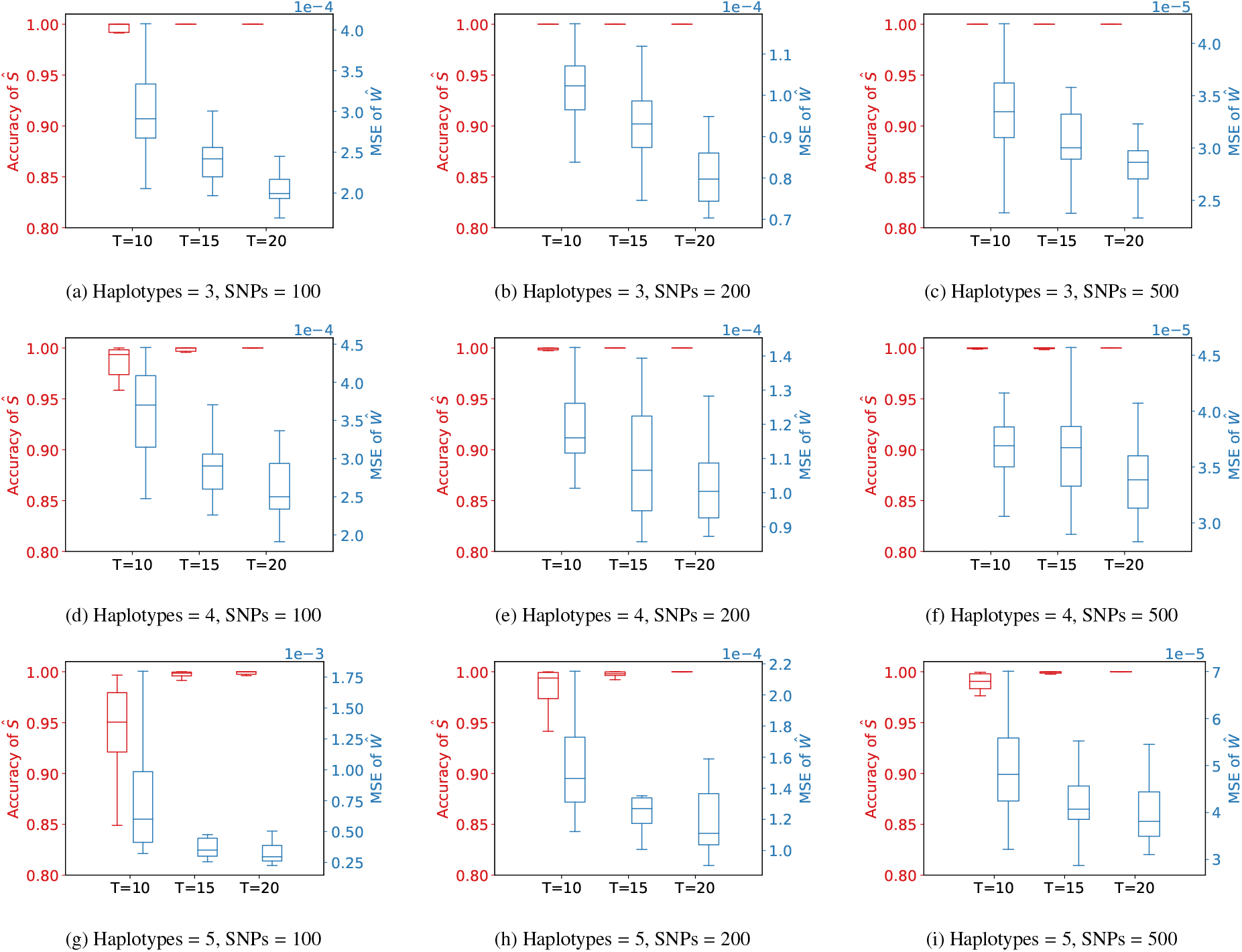
These plots are a set of boxplots to evaluate the performance of our estimation with different numbers m of haplotypes, numbers N of SNPs and numbers T of time points. Our noise level is *σ* = 0.05. The red boxplots present the accuracy rate of the estimated *S* matrix, which are calculated by the percentage of correctly estimated entries in posterior mean. And the blue boxplots demonstrate the MSE of the estimated *ω* matrix. Y axes are in the corresponding color. Each boxplot contains 20 groups of dataset. Accuracy rate and MSE are calculated by the posterior mean of 3000 steps in MCMC sampling excluding burn-in steps.

## References

[1] A. G. Clark, Inference of haplotypes from pcr-amplified samples of diploid populations., Molecular biology and evolution 7 (2) (1990) 111–122.

[2] B. L. Browning, S. R. Browning, A unified approach to genotype imputation and haplotype-phase inference for large data sets of trios and unrelated individuals, The American Journal of Human Genetics 84 (2) (2009) 210–223.

[3] Y. Li, C. J. Willer, J. Ding, P. Scheet, G. R. Abecasis, Mach: using sequence and genotype data to estimate haplotypes and unobserved genotypes, Genetic epidemiology 34 (8) (2010) 816–834.

[4] N. Barghi, R. Tobler, V. Nolte, A. M. Jakšić, F. Mallard, K. A. Otte, M. Dolezal, T. Taus, R. Kofler, C. Schlötterer, Genetic redundancy fuels polygenic adaptation in drosophila, PLoS biology 17 (2) (2019) e3000128.

[5] C. J. Illingworth, L. Parts, S. Schiffels, G. Liti, V. Mustonen, Quantifying selection acting on a complex trait using allele frequency time series data, Molecular biology and evolution 29 (4) (2012) 1187–1197.

[6] M. K. Burke, J. P. Dunham, P. Shahrestani, K. R. Thornton, M. R. Rose, A. D. Long, Genome-wide analysis of a long-term evolution experiment with drosophila, Nature 467 (7315) (2010) 587–590.

[7] A. Futschik, C. Schlotterer, The next generation of molecular markers from massively parallel sequencing of pooled dna samples, Genetics 186 (1) (2010) 207–218.

[8] C. Schlötterer, R. Tobler, R. Kofler, V. Nolte, Sequencing pools of individuals—mining genome-wide polymorphism data without big funding, Nature Reviews Genetics 15 (11) (2014) 749–763.

[9] S. U. Franssen, N. H. Barton, C. Schlötterer, Reconstruction of haplotype-blocks selected during experimental evolution, Molecular biology and evolution (2016) msw210.

[10] L. Excoffier, M. Slatkin, Maximum-likelihood estimation of molecular haplotype frequencies in a diploid population., Molecular biology and evolution 12 (5) (1995) 921–927.

[11] M. Pirinen, Estimating population haplotype frequencies from pooled snp data using incomplete database information, Bioinformatics 25 (24) (2009) 3296–3302.

[12] C.-C. Cao, X. Sun, Accurate estimation of haplotype frequency from pooled sequencing data and cost-effective identification of rare haplotype carriers by overlapping pool sequencing, Bioinformatics 31 (4) (2015) 515–522.

[13] D. Kessner, T. L. Turner, J. Novembre, Maximum likelihood estimation of frequencies of known haplotypes from pooled sequence data, Molecular biology and evolution 30 (5) (2013) 1145–1158.

[14] Q. Long, D. C. Jeffares, Q. Zhang, K. Ye, V. Nizhynska, Z. Ning, C. Tyler-Smith, M. Nordborg, Poolhap: inferring haplotype frequencies from pooled samples by next generation sequencing, PloS one 6 (1) (2011) e15292.

[15] M. Pelizzola, M. Behr, H. Li, A. Munk, A. Futschik, Multiple haplotype reconstruction from allele frequency data, Nature Computational Science 1 (4) (2021) 262–271.

[16] M. Behr, A. Munk, Statistical methods for minimax estimation in linear models with unknown design over finite alphabets, SIAM Journal on Mathematics of Data Science 4 (2) (2022) 490–513. arXiv:https://doi.org/10.1137/21M1398860, doi:10.1137/21M1398860. URL https://doi.org/10.1137/21M1398860

[17] S. Talwar, M. Viberg, A. Paulraj, Blind separation of synchronous co-channel digital signals using an antenna array. i. algorithms, IEEE Transactions on Signal Processing 44 (5) (1996) 1184–1197.

[18] A.-J. Van Der Veen, Asymptotic properties of the algebraic constant modulus algorithm, IEEE Transactions on Signal Processing 49 (8) (2001) 1796–1807.

[19] A. Leshem, N. Petrochilos, A.-J. van der Veen, Finite sample identifiability of multiple constant modulus sources, IEEE Transactions on Information Theory 49 (9) (2003) 2314–2319.

[20] Y. Zhang, S. A. Kassam, Blind separation and equalization using fractional sampling of digital communications signals, Signal Processing 81 (12) (2001) 2591–2608.

[21] C. Hans, Bayesian lasso regression, Biometrika 96 (4) (2009) 835–845.

[22] F. Caron, A. Doucet, Sparse bayesian nonparametric regression, in: Proceedings of the 25th international conference on Machine learning, 2008, pp. 88–95.

[23] N. G. Polson, J. G. Scott, Shrink globally, act locally: Sparse bayesian regularization and prediction, Bayesian statistics 9 (501-538) (2010) 105.

[24] R. Alhamzawi, K. Yu, D. F. Benoit, Bayesian adaptive lasso quantile regression, Statistical Modelling 12 (3) (2012) 279–297.

[25] T. Peltola, A. S. Havulinna, V. Salomaa, A. Vehtari, Hierarchical bayesian survival analysis and projective covariate selection in cardiovascular event risk prediction., BMA@ UAI 27 (2014) 79–88.

[26] X.-N. Feng, Y. Wang, B. Lu, X.-Y. Song, Bayesian regularized quantile structural equation models, Journal of Multivariate Analysis 154 (2017) 234–248.

[27] T. J. Mitchell, J. J. Beauchamp, Bayesian variable selection in linear regression, Journal of the american statistical association 83 (404) (1988) 1023–1032.

[28] I. M. Johnstone, B. W. Silverman, Needles and straw in haystacks: Empirical bayes estimates of possibly sparse sequences, The Annals of Statistics 32 (4) (2004) 1594–1649.

[29] M. Bogdan, A. Chakrabarti, F. Frommlet, J. K. Ghosh, Asymptotic bayes-optimality under sparsity of some multiple testing procedures, The Annals of Statistics 39 (3) (2011) 1551–1579.

[30] I. Castillo, A. van der Vaart, Needles and straw in a haystack: Posterior concentration for possibly sparse sequences, The Annals of Statistics 40 (4) (2012) 2069–2101.

[31] C. M. Carvalho, N. G. Polson, J. G. Scott, The horseshoe estimator for sparse signals, Biometrika 97 (2) (2010) 465–480.

[32] A. Armagan, M. Clyde, D. Dunson, Generalized beta mixtures of gaussians, Advances in neural information processing systems 24 (2011).

[33] A. Armagan, D. B. Dunson, J. Lee, Generalized double pareto shrinkage, Statistica Sinica 23 (1) (2013) 119.

[34] A. Bhattacharya, D. B. Dunson, Sparse Bayesian infinite factor models, Biometrika (2011) 291–306.

[35] M. Behr, A. Munk, Identifiability for blind source separation of multiple finite alphabet linear mixtures, IEEE Transactions on Information Theory 63 (9) (2017) 5506–5517.

[36] A. Gelman, D. Lee, J. Guo, Stan: A probabilistic programming language for bayesian inference and optimization, Journal of Educational and Behavioral Statistics 40 (5) (2015) 530–543.

[37] C. C. Monnahan, J. T. Thorson, T. A. Branch, Faster estimation of bayesian models in ecology using hamiltonian monte carlo, Methods in Ecology and Evolution 8 (3) (2017) 339–348.

[38] S. Relaxation, Gibbs distributions, and the bayesian restoration of images. s. geman and d. geman, IEEE Transactions on Pattern Analysis and Machine Intelligence 6 (1984) 721–741.

[39] M. Betancourt, Cruising the simplex: Hamiltonian monte carlo and the dirichlet distribution, arXiv preprint arXiv:1010.3436 (2010).

[40] F. Zanini, J. Brodin, L. Thebo, C. Lanz, G. Bratt, J. Albert, R. A. Neher, Population genomics of intrapatient hiv-1 evolution, Elife 4 (2015) e11282.

[41] L. M. Noble, M. V. Rockman, H. Teotónio, Gene-level quantitative trait mapping in caenorhabditis elegans, G3 11 (2) (2021) jkaa061.

